# Therapeutic targeting of SETD2-deficient cancer cells with the small-molecule compound RITA

**DOI:** 10.1101/2021.07.08.451582

**Authors:** Kirsten A. Lopez, Sovan Sarkar, Elena Seraia, Chiara Toffanin, Christian Cooper, Michalis Challoumas, Fiona A. Okonjo, George D. D. Jones, Francesca Buffa, Daniel Ebner, Timothy C. Humphrey

## Abstract

The histone methyltransferase SETD2 and its associated histone mark H3 lysine 36 trimethylation (H3K36me3) are frequently lost in certain cancer types, identifying SETD2 as an important therapeutic target. Here we show that SETD2-deficient cancer cells are profoundly sensitive to the compound RITA, resulting in significant p53 induction and apoptosis. This is further associated with defects in DNA replication, leading to delays in S-phase progression, increased recruitment of replication stress markers, and reduced replication fork progression. RITA sensitivity is linked to the phenol sulphotransferase SULT1A1, which we find to be highly upregulated in cells that lack SETD2. Depletion of SULT1A1 or addition of the phenol sulphotransferase inhibitor DCNP abolishes these phenotypes and suppresses the sensitivity of SETD2-deficient cancer cells, identifying SULT1A1 activity to be critical in mediating the potent cytotoxicity of RITA against SETD2-deficient cells. These findings define a novel therapeutic strategy for targeting the loss of SETD2 in cancer.

**Significance Statement:** The histone-modifying enzyme SETD2 has emerged as an important tumour suppressor in a number of different cancer types, identifying it as a promising therapeutic target. The concept of synthetic lethality, a genetic interaction in which the simultaneous loss of two genes or pathways that regulate a common essential process renders the cell nonviable, is a valuable tool for killing cancer cells that have known mutations. In this study, we conducted a synthetic lethality screen for compounds that specifically target SETD2-deficient cancer cells. The top hit, a compound called RITA, reduces cell viability and induces cell death only in the context of SETD2 loss, thereby highlighting a potential novel therapeutic strategy for treating SETD2-deficient cancers.

## Introduction

A key challenge in the treatment of cancer is achieving sufficient anti-tumour activity while limiting any negative impact on the patient. Maximising a drug’s therapeutic index, defined as the ratio between the dose that induces toxicity and the dose needed for therapeutic efficacy (1), is a problem that still plagues researchers and clinicians alike. This has led to increased efforts to discover tumour-specific therapies that can complement or even replace existing chemotherapies, which can induce severe side effects (2). A promising tool for achieving tumour specificity is the concept of synthetic lethality, a biological interaction in which two genes or pathways independently regulate the same cellular process (3).

Histone modifications are a critical mechanism through which cells control gene expression, chromatin structure, and genome stability. The dysregulation of the histone code is a common feature of tumourigenesis, with histone-modifying enzymes constituting some of the most frequently mutated genes in cancer (4–8). In mammals, SETD2 is the sole enzyme responsible for H3K36 trimethylation (H3K36me3) in somatic cells (9) and does so co-transcriptionally by binding to the phosphorylated C-terminal domain of RNA polymerase II (10–13). Consistently, H3K36me3 has been linked to active gene transcription (14) and loss of this histone mark is associated with the increased occurrence of cryptic intragenic transcription and dysregulated chromatin remodelling during gene transcription (15). H3K36me3 is also involved in the maintenance of genome stability via the regulation of the mismatch repair pathway (16) and homologous recombination (17). In addition, SETD2 has been demonstrated to methylate non-histone proteins such as microtubules, which prevents mitotic defects and aberrant chromosomal segregation (18).

The contribution of SETD2 to important genomic processes, including genome stability, is consistent with its role as a tumour suppressor in various cancer types. SETD2 deletions and mutations were detected in a substantial number of clear cell renal cell carcinoma (ccRCC) clinical samples (19) and cell lines (20). SETD2’s location on chromosome 3p makes it one of the most frequent examples of loss of heterozygosity (LOH) in ccRCC (21). Mutations in SETD2 are associated with lower cancer-specific survival in RCC (22), and analysis of H3K36me3 levels via immunohistochemistry revealed a progressive decline in staining intensity from primary tumour to metastases in RCC patients, culminating in an approximately 60% reduction in positive nuclei compared to uninvolved renal tissue (23). Missense or truncating mutations that inactivate the methyltransferase activity of SETD2 were also observed in 15% of paediatric and 8% of adult high-grade gliomas, with no mutations observed in tumours of Grade II and below (24). Comparison of SETD2 expression between breast tumour samples and adjacent normal tissue revealed a marked reduction in SETD2 transcript and protein levels in breast cancer (25).

Our group has previously reported an evolutionarily conserved synthetic lethal interaction between SETD2 and the cell cycle regulator WEE1 in fission yeast (26) and human cells (27). The WEE1 inhibitor AZD1775 was found to selectively target cancer cell lines that are SETD2-deficient and this coincided with the downregulation of RRM2, a subunit of the enzyme ribonucleotide reductase (RNR) (27). In order to further our understanding of SETD2’s role in tumourigenesis and identify complementary strategies for targeting SETD2 loss in cancer, we conducted a high-throughput small-molecule compound screen. More than 2000 compounds were tested on U2OS osteosarcoma cells in which the SETD2 gene has been deleted using the CRISPR/Cas9 system, as well as on isogenic parental U2OS cells. The compound that showed the most drastic anti-tumour activity specifically against SETD2-CRISPR cells was RITA (**R**eactivation of p53 and **I**nduction of **T**umour cell **A**poptosis), which was initially discovered due to its high potency and selectivity against a number of tumour types in the National Cancer Institute (NCI) Anticancer Drug Screen, which used a cell line panel comprised of 60 different human cancer cell lines (28). RITA has been reported to bind p53, prevent its interaction with MDM2, and restore its functions in transactivation and apoptosis (Issaeva et al., 2004). Here we report our findings that RITA potently targets SETD2-deficient cancer cells, leading to strikingly reduced cell viability and increased apoptosis associated with p53 upregulation and activation, activation of the DNA damage response, and disruption of DNA replication.

## Results

### SETD2-deficient cancer cells are hypersensitive to RITA

In order to discover novel therapeutic strategies for targeting the loss of H3K36me3 in cancer, a high-throughput small-molecule compound synthetic lethality screen was conducted. More than 2000 compounds were tested on CRISPR/Cas9 SETD2-deleted U2OS osteosarcoma cells (27), as well as on isogenic parental U2OS cells. The details of the compound libraries used are in Table S1.

After performing statistical analysis and ranking the hits based on their Z-scores, the compound that showed the strongest anti-tumour activity specifically against SETD2-CRISPR cells was RITA (**R**eactivation of p53 and **I**nduction of **T**umour cell **A**poptosis), also known as NSC 652287. Initial doseresponse experiments of up to 20 μM showed a striking difference in sensitivity between parental and SETD2-CRISPR cells (Fig. S1), and this difference was maintained at nanomolar dose ranges (Fig. 1A). RITA was initially discovered due to its high potency and selectivity against a number of tumour types in the National Cancer Institute (NCI) Anticancer Drug Screen, (28). Sensitivity was reported to be particularly striking in several renal cell carcinoma (RCC) cell lines, including the A498 cell line, in which SETD2 is mutated. To confirm whether this result could be replicated in our system, dose-response curves for RITA were obtained in the RCC cell lines A498, LB996, and 786-O. Even at low concentrations of RITA, both SETD2-mutant cell lines (A498 and LB996) were markedly more sensitive compared to the SETD2-wild type cell line 786-O (Fig. 1B). The specific cytotoxicity of RITA against SETD2-deficient cells was further assessed via clonogenic survival assays of SETD2 wild-type versus CRISPR cells. SETD2-deficient cells exhibited exquisite sensitivity to RITA, thus corroborating the results of the previous viability assays (Fig. 1C). Together, these observations indicate that SETD2 loss can be specifically and potently targeted by using the compound RITA.

**Figure 1.**
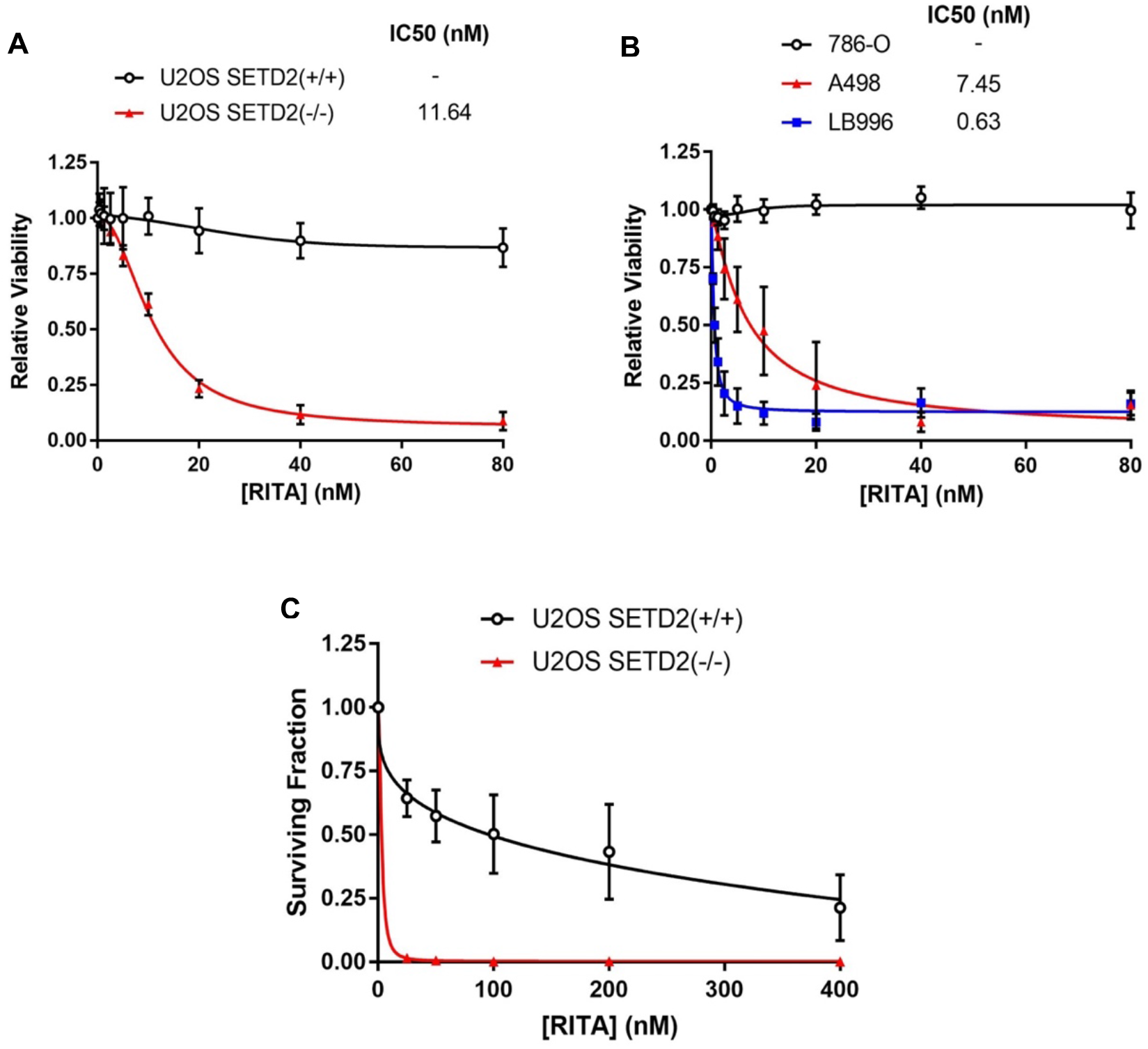
SETD2-deficient cells are hypersensitive to RITA. **(A-B)** Dose response viability curves for **(A)** U2OS cells and **(B)** renal cell carcinoma cell lines with wild-type (786-O) or mutant (A498 and LB996) SETD2 treated with RITA. Data are shown as mean ± SD. **(C)** Clonogenic survival assay for parental and SETD2-CRISPR U2OS treated with RITA. Data are shown as mean ± SD.

### RITA induces p53 upregulation and activation, as well as apoptotic cell death, specifically in SETD2-deficient cancer cells

One possible mechanism underlying RITA’s remarkable potency may involve its ability to stabilize the p53 protein. RITA has been reported to bind p53, prevent its interaction with MDM2, and restore its functions in transactivation and apoptosis (29). Furthermore, p53 has been shown to interact with SETD2, which is capable of regulating certain p53 target genes, including MDM2 (30). Consistent with published reports in other cell lines, RITA treatment led to a striking increase in p53 protein levels in SETD2-CRISPR U2OS and SETD2-mutant A498 cells (Fig. 2A). In addition, RITA greatly induced the phosphorylation of p53 at Ser15 specifically in SETD2-deficient U2OS and A498 cells (Fig. 2A).

**Figure 2.**
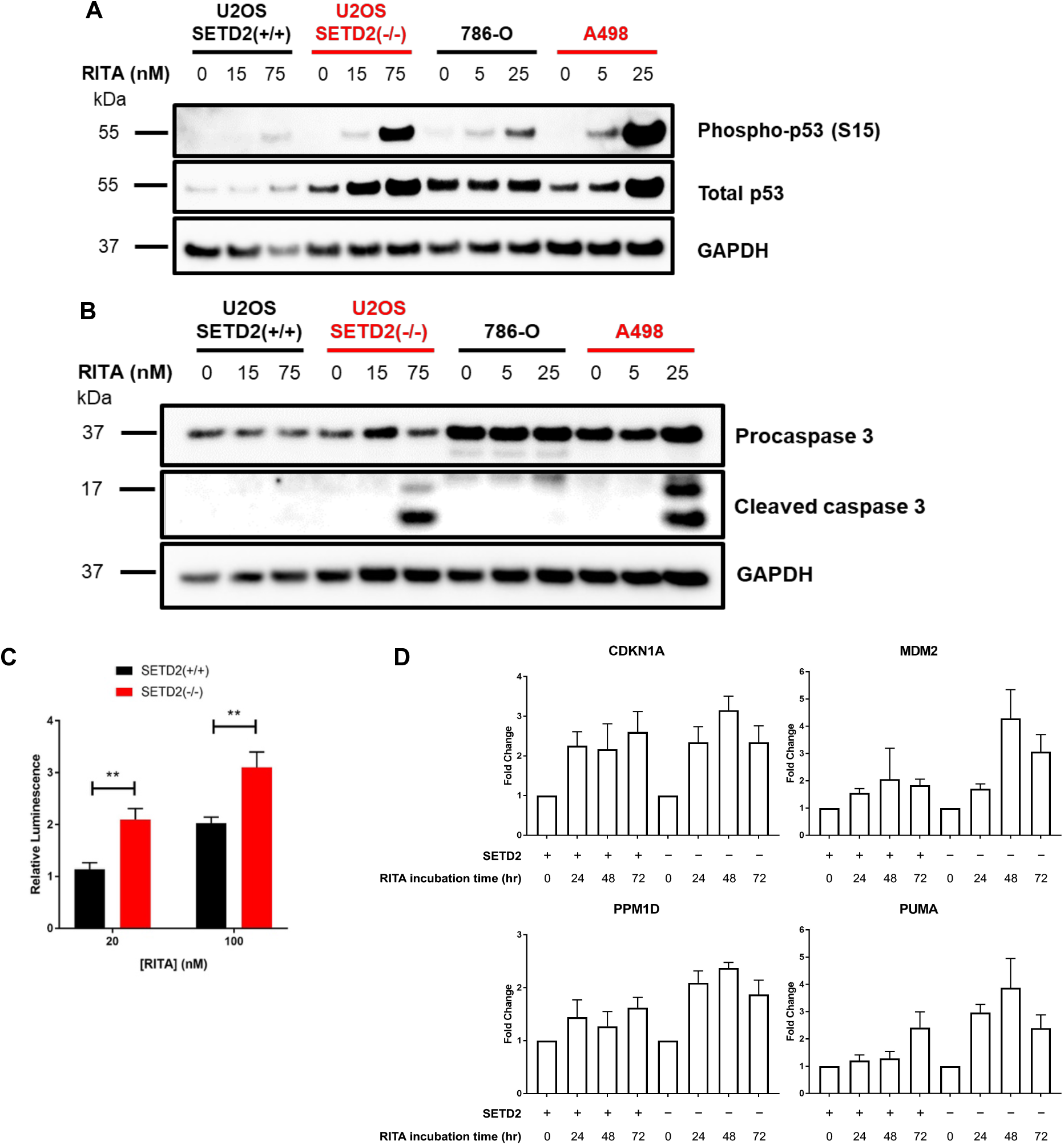
RITA stabilizes and activates p53 and induces apoptosis in SETD2-deficient cancer cells. **(A)** Western blot of total and phospho-p53 (Ser15) in parental and SETD2-CRISPR U2OS and RCC cell lines treated with the indicated doses of RITA for 48 hours. **(B)** Western blot of procaspase 3 and cleaved caspase 3 in SETD2-CRISPR and parental U2OS and RCC cell lines after RITA treatment for 48 hours. **(C)** Annexin V levels in SETD2-CRISPR and parental U2OS cells after RITA treatment for 30 hours. Luminescence readings were normalized to untreated controls. Data are shown as mean ± SD. P-values were calculated using the two-tailed Student’s t-test. **p < 0.01. **(D)** Quantitative RT-PCR analysis of p53 target gene expression after RITA treatment. Samples were normalised to the housekeeping gene *GAPDH*. Fold change was calculated relative to untreated controls. Data are shown as mean ± SD. **(E)** Quantitative RT-PCR analysis of *TP53* and its target genes in untreated SETD2-CRISPR U2OS. Fold change was calculated relative to untreated parental U2OS.

Given p53’s established role in promoting apoptotic cell death in response to radiation and chemotherapy (31, 32), the induction of apoptosis was measured after exposure to RITA. Western blot analysis of caspase-3 cleavage revealed elevated levels in SETD2-CRISPR U2OS and A498 cells, but not their SETD2-wildtype counterparts, in the presence of RITA (Fig. 2B). Using a luminescence-based assay, we also found annexin V levels to be significantly elevated in SETD2-CRISPR U2OS cells after RITA treatment (Fig. 2C), indicating that apoptotic cell death was greatly induced in these cells in response to RITA.

To verify whether the observed upregulation of p53 was leading to activation of its downstream targets, the expression levels of some canonical p53 target genes were analysed by quantitative RT-PCR. Indeed, mRNA levels of the genes CDKN1A, MDM2, PPM1D, and PUMA were increased after RITA treatment, and this effect was stronger in the absence of SETD2 (Fig. 2D). In the case of CDKN1A, RITA induced gene expression to a similar extent in both parental and SETD2-CRISPR cells, but it is worth noting that CDKN1A expression is already more than 2-fold higher after SETD2 deletion in unperturbed conditions (Fig. 2E); thus, CDKN1A expression after RITA treatment is still 2-fold higher in SETD2-CRISPR cells relative to parental cells. Notably, p53 mRNA levels were not substantially affected by RITA treatment in SETD2-CRISPR cells (Fig. S2), indicating that the striking increase in protein expression is mediated by post-translational mechanisms. Overall, these results demonstrate that RITA leads to the upregulation and activation of p53, associated with the induction of p53-mediated transcription and apoptosis, particularly in cells that lack functional SETD2.

### RITA induces replication stress, the DNA damage response and cell cycle arrest specifically in SETD2-deficient cancer cells

To investigate the mechanism through which RITA targets SETD2-deficient cells, we looked at the effect of RITA treatment on cell cycle progression. In both SETD2-CRISPR U2OS and SETD2-mutant A498 cells, RITA induced a dose-dependent accumulation in non-replicating S-phase (defined as having a DNA content between 2N and 4N but no incorporation of the synthetic nucleoside BrdU) and G2/M (Fig. 3A, Fig. S3). These results suggest a disruption in DNA replication and the activation of the DNA damage response, leading to the activation of cell cycle checkpoints. We reported a similar phenotype upon treatment of SETD2-deficient cells with the WEE1 inhibitor AZD1775, which was shown to be mediated by downregulation of the ribonucleotide reductase subunit RRM2 and subsequent disruption of the cellular dNTP supply (27). To determine whether the same mechanism is involved with RITA, we looked at whether RITA sensitivity can be modulated by the addition of exogenous dNTPs. Viability assays in both parental and SETD2-CRISPR U2OS cells demonstrated that supplementation with additional nucleosides has no effect on RITA’s potency (Fig. S4A). These results were further corroborated by the finding that RRM2 protein levels, while reduced in the absence of SETD2 as previously reported (27), do not change after RITA treatment (Fig. S4B). Thus, there is currently no evidence to support a role for dNTP maintenance in the sensitivity of SETD2-deficient cells to RITA.

**Figure 3.**
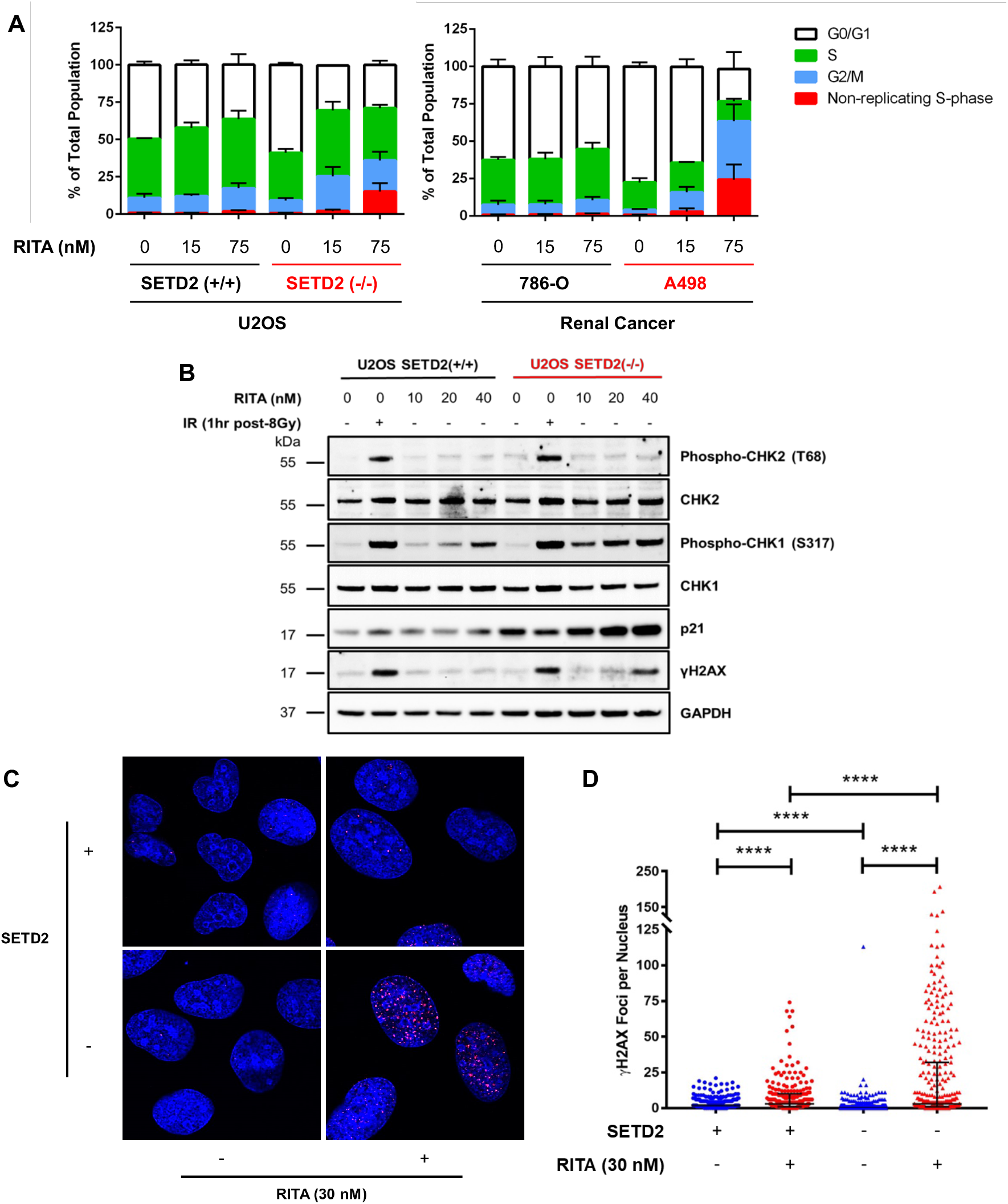
RITA induces cell cycle arrest and activates the DNA damage response in SETD2-deficient cancer cells. **(A)** Cell cycle profiles of parental and SETD2-CRISPR U2OS and RCC cell lines after RITA treatment. Data shown as mean ± SD. **(B)** Western blot of total and phospho-CHK1 (Ser317), total and phospho-CHK2 (Thr68), p21, and γH2AX in parental and SETD2-CRISPR U2OS after RITA treatment. **(C)** Confocal microscopy of γH2AX foci (red) in parental and SETD2-CRISPR U2OS cells after RITA treatment. Representative images from 3 independent experiments are shown. **(D)** Quantification of γH2AX foci per nucleus from **(C)**. At least 100 nuclei per biological replicate (n = 3) were counted for statistical analysis. Data shown as median + interquartile range. P-values were calculated using the two-tailed Student’s t-test.

Consistent with the observed cell cycle defects, detection of key checkpoint mediators by Western blotting revealed the activation of CHK1, phosphorylation of histone H2AX (γH2AX), and upregulation of the CDK inhibitor p21 (CDKN1A) by RITA, which occurred to a markedly stronger degree in SETD2-CRISPR cells compared to their parental counterparts (Fig. 3B). In line with its mRNA expression levels (Fig. 2D-E), p21 is much more highly expressed in SETD2-CRISPR cells and is further upregulated upon RITA treatment (Fig. 3B). The presence of DNA damage after RITA treatment, specifically in the absence of SETD2, was further verified by visualising γH2AX foci by immunofluorescence microscopy (Fig. 3C-D). Interestingly, there was no evidence of CHK2 activation in the presence of RITA, suggesting that RITA-induced DNA damage does not lead to the formation of DNA double-strand breaks (DSBs). Therefore, these findings indicate that, in the absence of functional SETD2, RITA treatment leads to some form of DNA damage or genotoxic stress that activates the DNA damage response specifically via the ATR-CHK1 axis, and this is associated with impaired DNA synthesis and cell cycle arrest at the G2/M checkpoint.

One of the key questions in elucidating RITA’s mechanism of action in the context of SETD2 deficiency is the importance of SETD2 and H3K36me3’s myriad of cellular functions. Most pertinent of these to the phenotypes observed thus far is the published role of H3K36me3 in the repair of DSBs via homologous recombination (HR) (17), which is consistent with the increased DDR activation, replication stress, and cell death induced by RITA in the absence of SETD2. If HR plays a substantial role in the RITA sensitivity of SETD2-deficient cells, we can hypothesise that other HR factors would behave similarly in this context. However, siRNA-mediated knockdown of RAD51 and BRCA1 in SETD2-positive U2OS cells and 786-O cells did not sensitize them to RITA to the same extent as their SETD2-deficient counterparts, even at high drug concentrations above 1 μM (Fig. S4C). Furthermore, in U2OS cells, no dose-dependent increase in RITA sensitivity was observed after depletion of RAD51 or BRCA1. Therefore, these results suggest that SETD2’s function in the repair of DNA damage via homologous recombination is not the main mechanism through which RITA exerts its cytotoxicity.

The absence of CHK2 phosphorylation implies that RITA is inducing replication stress and not DSBs (Fig. 3B). To determine whether DNA replication is perturbed after RITA treatment, parental and SETD2-CRISPR U2OS cells were stained for the replication stress markers phospho-RPA (Ser33) and 53BP1. RITA treatment led to significantly higher levels of both types of foci particularly in SETD2-CRISPR cells (Fig. 4A-B), indicating increased replication stress in the presence of RITA. Furthermore, SETD2-CRISPR cells displayed a significant reduction in replication fork velocity after RITA treatment, whereas the parental cells remained unaffected (Fig. 4C-D). Moreover, this was observed within a short treatment window (30 minutes), indicating that RITA enters the cell and interferes with DNA replication very rapidly. Overall, these results demonstrate that RITA activates the DNA damage response by rapidly impairing DNA replication fork progression and inducing replication stress, as demonstrated by increased phosphorylation of RPA and recruitment of 53BP1.

**Figure 4.**
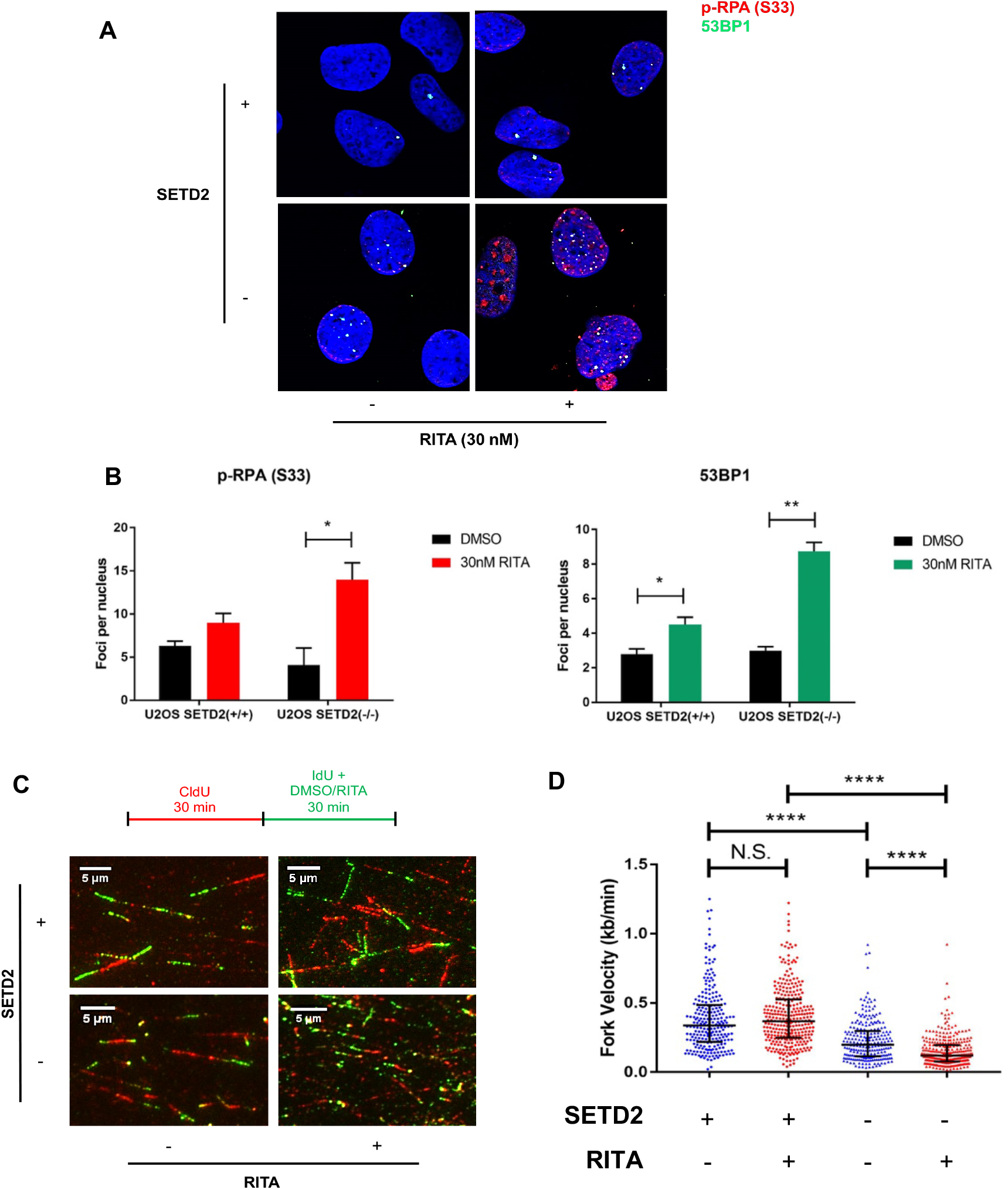
RITA induces replication stress in SETD2-deficient cancer cells. **(A)** Confocal microscopy of phospho-RPA Ser33 (red) and 53BP1 (green) foci in U2OS cells after RITA treatment. Representative images from 3 independent experiments are shown. **(B)** Quantification of the average number of foci per nucleus from **(A)**. At least 70 nuclei per biological replicate (n = 3) were counted for statistical analysis. Data shown as mean ± SD. P-values were calculated using the two-tailed Student’s t-test. *p < 0.05, **p < 0.01. RITA induces replication stress in SETD2-deficient cancer cells. **(C)** DNA fibre assay of parental and SETD2-CRISPR U2OS. Cells were treated as indicated. Representative images from 2 independent experiments are shown. **(D)** Quantification of replication fork velocity from **(C)**. At least 200 fibres per biological replicate (n = 3) were measured for statistical analysis. Data shown as median + interquartile range. P-values were calculated using the two-tailed Student’s t-test. N.S. = non-significant, ****p < 0.000001.

RITA has been previously reported to act directly on DNA via crosslinking (33). The presence of RITA-induced replication stress in SETD2-deficient cells supports this notion, as crosslinking agents have been shown to mediate cellular cytotoxicity in a replication-dependent manner (34). To test this hypothesis, we directly measured crosslink formation via a modified comet assay. Whereas the positive control, the crosslinking agent mitomycin C, showed a striking reduction in fragmented (tail) DNA after IR, we did not observe any such decrease after RITA treatment (Fig. S5A). This was further corroborated by the lack of sensitivity of SETD2-CRISPR cells to mitomycin C (Fig. S5B). Given the key role played by the HR pathway in ICL repair (35), the absence of crosslinks in this case is consistent with the minimal effects of HR deficiency on RITA sensitivity (Fig. S4C).

### RITA sensitivity is mediated via enhanced SULT1A1 sulphotransferase activity in SETD2-deficient cells

Despite initial reports claiming that RITA’s cytotoxicity is mainly driven by its ability to reactivate p53 (29), there have been a number of subsequent findings that link RITA’s anticancer activity to p53-independent mechanisms (36–39). In particular, a significant correlation between RITA sensitivity and expression of the phenol sulphotransferase SULT1A1 was observed in renal cell carcinoma cell lines (40). To determine whether SULT1A1 plays a role in the interaction between SETD2 loss and RITA, we performed siRNA depletion of SULT1A1 in SETD2-CRISPR U2OS cells. The addition of SULT1A1-specific siRNA prior to treatment significantly modulated the RITA sensitivity of SETD2(-/-) U2OS cells compared to control siRNA (Fig. 5A). In accordance with published reports, SULT1A1 protein (Fig. 5B) and mRNA (Fig. S6A) levels were strikingly upregulated in RITA-sensitive SETD2-deleted U2OS cells compared to their parental counterparts. These results suggest a key role for SULT1A1 in mediating the potent cytotoxicity of RITA against SETD2-deficient cancer cells.

**Figure 5.**
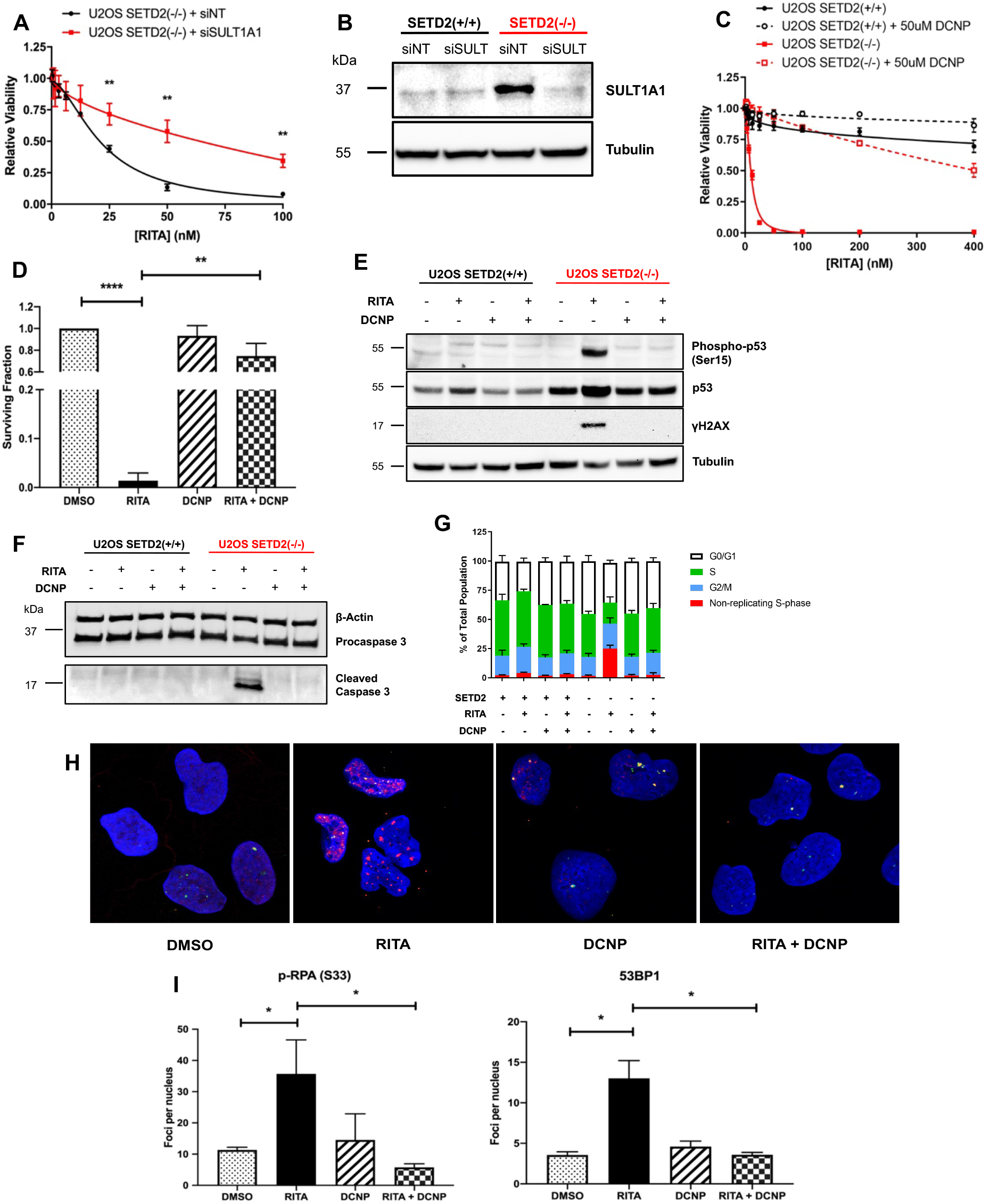
Loss or inhibition of the phenol sulphotransferase SULT1A1 abolishes RITA-induced phenotypes. **(A)** Dose response viability curves for SETD2-CRISPR U2OS cells treated with RITA after the addition of non-targeting control (siNT) or SULT1A1 siRNA (siSULT1A1). Data are shown as mean ± SD. **(B)** Western blot of SULT1A1 in U2OS cells in the presence of non-targeting control (siNT) or SULT1A1 siRNA (siSULT). **(C)** Dose response viability curves for U2OS cells treated with RITA in the presence or absence of DCNP. Data are shown as mean ± SD. **(D)** Clonogenic survival assay for U2OS cells in the presence of the indicated compounds. Data are shown as mean ± SD. P-values were calculated using the two-tailed Student’s t-test. **p < 0.01, ****p < 0.0001. **(E)** Western blot of total p53, phospho-p53 (Ser15), and γH2AX in U2OS cells after the indicated treatments. **(F)** Western blot of procaspase 3 and cleaved caspase 3 in U2OS cells after the indicated treatments. **(G)** Cell cycle profiles of U2OS cells in the presence of the indicated compounds. Data shown as mean ± SD. **(H)** Confocal microscopy of phospho-RPA Ser33 (red) and 53BP1 (green) foci in U2OS cells after the indicated treatments. Representative images from 3 independent experiments are shown. **(I)** Quantification of the average number of foci per nucleus from **(I)**. At least 80 nuclei per biological replicate (n = 3) were counted for statistical analysis. Data shown as mean ± SD. P-values were calculated using the two-tailed Student’s t-test. *p < 0.05.

To further confirm the involvement of SULT1A1, we tested the phenol sulphotransferase inhibitor 2,6-dichloro-4-nitrophenol (DCNP) (41) in combination experiments with RITA. DCNP on its own displayed minimal toxicity against U2OS cells regardless of SETD2 status at concentrations of up to 100 μM (Fig. S6B). The presence of DCNP significantly reduced the RITA sensitivity of SETD2-CRISPR U2OS cells (Fig. 5C), rescuing viability to levels comparable to that of parental U2OS. We observed a similar impact on clonogenic survival (Fig. 5D), which showed the DCNP/RITA combination having almost the same surviving fraction as the DMSO control. These findings support the hypothesis that RITA requires SULT1A1 activity to exert its cytotoxic effects on cells lacking SETD2.

We next investigated whether SULT1A1 inhibition modulated other RITA-induced phenotypes. The levels of total and Ser15-phosphorylated p53, which were strikingly upregulated upon RITA treatment, were reverted to basal levels in the presence of DCNP (Fig. 5E). Consistently, the occurrence of apoptotic cell death as measured by cleaved caspase-3 levels was also suppressed by DCNP (Fig. 5F). These results suggest that RITA’s intracellular effects are dependent on SULT1A1 activity. Furthermore, FACS analysis of cell cycle progression showed a clear reduction in the non-replicating S-phase population when DCNP is added together with RITA (Fig. 5G), which indicates that SULT1A1 is necessary for the replication defects that occur after RITA treatment. In support of this, the elevated recruitment of replication stress markers phospho-RPA (Ser33) and 53BP1 in the presence of RITA was abolished by DCNP co-treatment (Fig. 5G-I). Overall, these data indicate that the potent and specific targeting of SETD2-deficient cells by RITA is dependent on the activity of the phenol sulphotransferase SULT1A1.

## Discussion

Rapid and continuous improvements in biotechnology, especially in genome sequencing, have been accompanied by a rise in personalised medicine, where a patient’s genotype informs his or her therapeutic options to maximise clinical benefits and minimise adverse effects. In oncology, personalised or precision medicine is enabled by the concept of synthetic lethality and has been demonstrated repeatedly over the years, most prominently in the case of BRCA mutations and PARP inhibition (42, 43). The aim of this study was to apply this idea to SETD2-deficient cancers in order to provide new and complementary strategies in addition to our published report on the SETD2-WEE1 synthetic lethality (27). In a large-scale screen comprised of thousands of compounds, the hit that displayed the highest potency against SETD2-deleted cells by a significant margin was RITA. In some ways, the phenotypes induced by RITA in SETD2-deficient cells reflect the observations made with SETD2 and WEE1. The most obvious of these is the presence of replication stress, as denoted by the increase in non-replicating S-phase (Fig. 3A), the activation of CHK1 (Fig. 3B), the accumulation of phospho-RPA and 53BP1 foci (Fig. 4A-B), and the decrease in replication fork velocity (Fig. 4C-D) after RITA treatment in the absence of SETD2. However, RITA is >10-fold more potent in reducing the viability of SETD2-deficient cells compared to the WEE1 inhibitor AZD1775, based on published data for the latter (11.6 nM vs. 151 nM for U2OS SETD2-CRISPR, 7.5 nM vs. 87 nM for A498, and 0.6 nM vs 68 nM for LB996) (Fig. 1A-B) (27). There are, however, key differences in the mechanisms of action between RITA and AZD1775. Despite the critical role of RRM2 and dNTP homeostasis in the SETD2-WEE1 synthetic lethality, there is no indication that bolstering the dNTP supply of SETD2-deficient cells has any significant effect on RITA sensitivity nor does RITA have any impact on RRM2 levels (Fig. S4A-B). Furthermore, we observed that homologous recombination, despite being one of the critical functions of this histone mark (17, 27), is not a major factor in RITA’s cytotoxicity against SETD2-deficient cells (Fig. S4C), which is consistent with the absence of a DSB response via ATM/CHK2 (Fig. 3B).

The molecular mechanisms that RITA employs against a variety of tumours and cell lines have been the subject of many studies. Although most frequently associated with its inhibition of the p53-MDM2 interaction (29), a number of groups have reported RITA cytotoxicity in p53-independent settings. This has been attributed to several different mediators, such as mitogen-activated protein kinases (MAPK) p38 and JNK (36), the Hedgehog pathway (37), and pro-survival and anti-apoptotic pathways distinct from those regulated by p53 (38). Perhaps most contradictory to the notion of RITA’s dependence on p53 is a CRISPR screen conducted to shed light on this issue; whereas nutlin-3 treatment led to a significant enrichment in cells with CRISPR-induced insertion/deletion (indel) mutations or truncations in the *TP53* locus, RITA treatment did not (39). Our results show unequivocally that RITA induces p53 upregulation and activation in the absence of SETD2 (Fig. 2A), associated with increased caspase-3 cleavage (Fig. 2B) and apoptotic cell death (Fig. 2C). It is possible that the impact on p53 is a consequence and not a cause of RITA’s intracellular activities; we observed that p53-CRISPR cells are less sensitive to RITA than their isogenic counterparts but siRNA-mediated knockdown of SETD2 was able to enhance RITA’s detrimental effects on cell viability regardless of p53 status (Fig. S7A-B).

Much of the uncertainty regarding RITA’s biological effects stems from conflicting evidence about its molecular target. One potential candidate, the phenol sulphotransferase SULT1A1, was described in a study that used transcriptomic profiling to predict sensitivity to almost 500 compounds, which calculated a significant correlation between RITA sensitivity and SULT1A1 mRNA and protein expression (40). Reducing SULT1A1 activity by using either siRNA-mediated depletion or a phenol sulphotransferase inhibitor corroborates these published reports and demonstrates that modulating this enzyme impairs RITA’s cytotoxicity in the absence of SETD2. This is also consistent with existing literature that indicates SULT1A1 is required to activate other carbinol-based compounds similar to RITA (44). Interestingly, the same report suggests that p53 loss is associated with SULT1A1 loss (44), which may explain why p53 deletion in some cell lines such as HCT116 abrogates RITA sensitivity. Our observation that SETD2-CRISPR cells have both high p53 and high SULT1A1 expression points to a mechanistic link between these genes, one that necessitates further investigation.

In summary, we have demonstrated a novel synthetic lethal interaction between SETD2 and the compound RITA. RITA potently and specifically targets SETD2-deficient cells and upregulates p53, leading to activation of downstream target genes and apoptotic cell death. RITA induces non-replicating S-phase and G2/M arrest in SETD2-deficient cells and activates the DNA damage response, leading to increased levels of phospho-CHK1, p21, and γH2AX. This is further associated with increased recruitment of replication stress markers phospho-RPA and 53BP1 and reduced replication fork velocity, suggesting a disruption in the process of DNA replication. Depletion or inhibition of the phenol sulphotransferase SULT1A1 abrogates these phenotypes and both p53 and SULT1A1 are highly upregulated in the absence of SETD2. This implies a feedback mechanism that links these two tumour suppressors with an enzyme that, while necessary for the production of steroid hormones and detoxification of xenobiotics, may inadvertently enhance the activity of cytotoxic carbinol compounds.

## Materials and Methods

### Cell Lines

The U2OS cell line was derived from human osteosarcoma and obtained from ATCC (ATCC number HTB-96). The U2OS SETD2-deleted cell line was generated by our own group using the CRISPR-Cas9 technology (27).

The 786-O, A498, and LB996 cell lines were derived from human clear cell renal cell carcinomas. 786-O was obtained from the lab of Dr. Valentine Macaulay at the Department of Oncology, University of Oxford. A498 was obtained from ATCC (HTB-44). LB996 was obtained from the lab of Prof. Benoit van den Eynde at the Oxford Ludwig Institute.

### Cell Culture

All cell lines except LB996 were maintained in complete medium consisting of Dulbecco’s modified minimal essential medium (DMEM, Life Technologies) supplemented with 10% foetal bovine serum (FBS, Sigma), 100 U penicillin (Sigma), and 0.1 mg/ml streptomycin (Sigma). LB996 cells were cultured in Iscove’s Modified Dulbecco’s Medium (IMDM, Life Technologies) with 10% FBS, 100 U penicillin, 0.1 mg/ml streptomycin, and G5 supplement (Life Technologies). All cells were grown at 37°C in a humidified incubator with 5% carbon dioxide. Cells were subcultured every 3-4 days as follows: cells were washed with phosphate buffered saline (PBS, Life Technologies) and then incubated with trypsin (Life Technologies) at 37°C until most cells had detached. Trypsinised cells were re-suspended in complete medium, centrifuged at 300 *g* for 3 minutes, and re-plated at the appropriate dilution.

### High-throughput Compound Screening

Parental and SETD2-deleted U2OS cells were seeded into 384-well plates (750 cells in 75 μl per well) using the Janus Liquid Handling Workstation (Perkin Elmer) and incubated at 37°C overnight. Subsequently, the compound libraries (TDI Expanded Oncology Drug Set and SelleckChem Bioactive Compound Library, 10 mM in DMSO) were thawed at room temperature and serially diluted into complete DMEM containing dimethyl sulphoxide (DMSO) using the Echo Acoustic Liquid Handler (Labcyte). The diluted compounds (or DMSO as a negative control) were added to the cells (75 μl per well) using the Janus workstation and incubated for 24 hours. The following day, the WEE1 inhibitor AZD1775 was diluted and added to the cells using the same procedure as the compound libraries and incubated for 72 hours. Subsequently, cell viability was measured using the resazurin assay.

### Statistical Analysis

Statistical analysis of the compound screening results was performed using the HTScape software, which was developed in-house by Dr. Francesca Buffa’s group. The raw data (.csv) files obtained from the resazurin assay and annotation files for the compound libraries were uploaded onto the HTScape software and Z-scores were calculated. Comparison between different cell lines and conditions was carried out by subtracting their respective Z-scores and calculating a false-positive discovery rate (PFP). Hits with PFP < 0.05 were defined as statistically significant.

Statistical analysis of other experimental results was performed using the GraphPad Prism 8 software.

### Validation of Positive Hits

2000 cells/well were seeded into a 96-well plate overnight before drug/inhibitor treatment. DMSO (solvent for the inhibitors) was used as a negative control. Cells were first treated with either DMSO or 2 μM RO3306, a CDK1 inhibitor, for 24 hours. This was followed by the addition of increasing concentrations (from a maximum of 20 μM serially diluted in increments of 2) of the compound to be validated on top of the existing medium, after which the cells were incubated for another 48 hours prior to measurement of cell viability via the resazurin assay.

### Drugs and Inhibitors

RITA (NSC 652287) and 2,6-dichloro-4-nitrophenol (DCNP) were obtained from Selleck Chemicals. All inhibitors were dissolved in DMSO and stored at −80°C.

### Resazurin Assay for Cell Viability

At the desired time point after treatment, culture medium was removed and 100 μl of fresh complete medium without phenol red and containing 20 μg/ml resazurin (Sigma) was added to each well. Resazurin is a nonfluorescent dye that can be converted via a redox reaction to a red fluorescent compound, resorufin, by living cells. The fluorescent signal is proportional to the number of living cells and was measured by a fluorescence plate reader (BMG Labtech) after 2 hours of incubation at 37°C. For drug treatment experiments, IC50 is defined as the concentration of drug that reduces viability by 50%.

### Clonogenic Survival Assay

Prior to drug treatment, cells were seeded into 6-well plates (2mL per well) at low densities and incubated overnight at 37°C. Culture media was subsequently replaced with media containing DMSO or the appropriate concentration of drug and incubated for 10 days at 37°C. Each well was then washed with PBS and stained with Brilliant Blue R Concentrate (Sigma) for 1 hour with gentle shaking. Plating efficiency (PE) and Surviving Fraction (SF) were calculated according to the published protocol (45).

### Apoptosis Detection Assay

Apoptotic cell death was measured in a 96-well format using the RealTime-Glo™ Annexin V Apoptosis and Necrosis Assay (Promega) per the manufacturer’s instructions. In brief, cells were seeded at 3000 cells per well (100 μl per well) in 96-well plates overnight. The following day, the media was replaced with 100 μl/well of media containing the compound to be tested. Immediately afterwards, another 100 μl/well containing the 2X Detection Reagent (which consists of Annexin NanoBiT® Substrate, CaCl_2_, Necrosis Detection Reagent, Annexin V-SmBiT, and Annexin V-LgBiT) was added. The luminescence signal is proportional to the number of apoptotic cells and was measured by a plate reader (BMG Labtech) 30 hours after incubation at 37°C.

### siRNA Transfection

All siRNAs (10-20 nM final concentration) were delivered to the cells by Lipofectamine RNAiMAX (Life Technologies) per the manufacturer’s instructions. Cells were incubated with siRNA-Lipofectamine mix in suspension (reverse transfection) in complete medium without penicillin and streptomycin. The medium was replaced 16 hours after transfection. Cells were analysed or further treated at least 48 hours after transfection. Western blotting was performed to determine knockdown efficiency. The sequences of the siRNAs are listed below:

siSETD2 (pooled) (Dharmacon): GAAACCGUCUCCAGUCUGU, UAAAGGAGGUAUAUCGAAU
siSULT1A1 (pooled) (Life Technologies): ACCAAGCGGCUCAAGAAUAAA, GAGAAGUUCAUGGUCGGAGAA

### Quantitative RT-PCR

Total RNA was purified using the RNeasy kit (Qiagen) and cDNA was prepared using the SuperScript RT-PCR system (Invitrogen). Quantitative RT-PCR was performed using the SYBR™ Green PCR Master Mix (Applied Biosystems) according to the manufacturer’s protocol. Reactions were carried out in triplicates for each target transcript using a 7500 Fast Real-Time PCR System (Applied Biosystems). The comparative C_T_ method was applied for quantifying the gene expression, and values were normalised against GAPDH as a control. Results were expressed as fold changes compared to the control condition. The following primers were used:

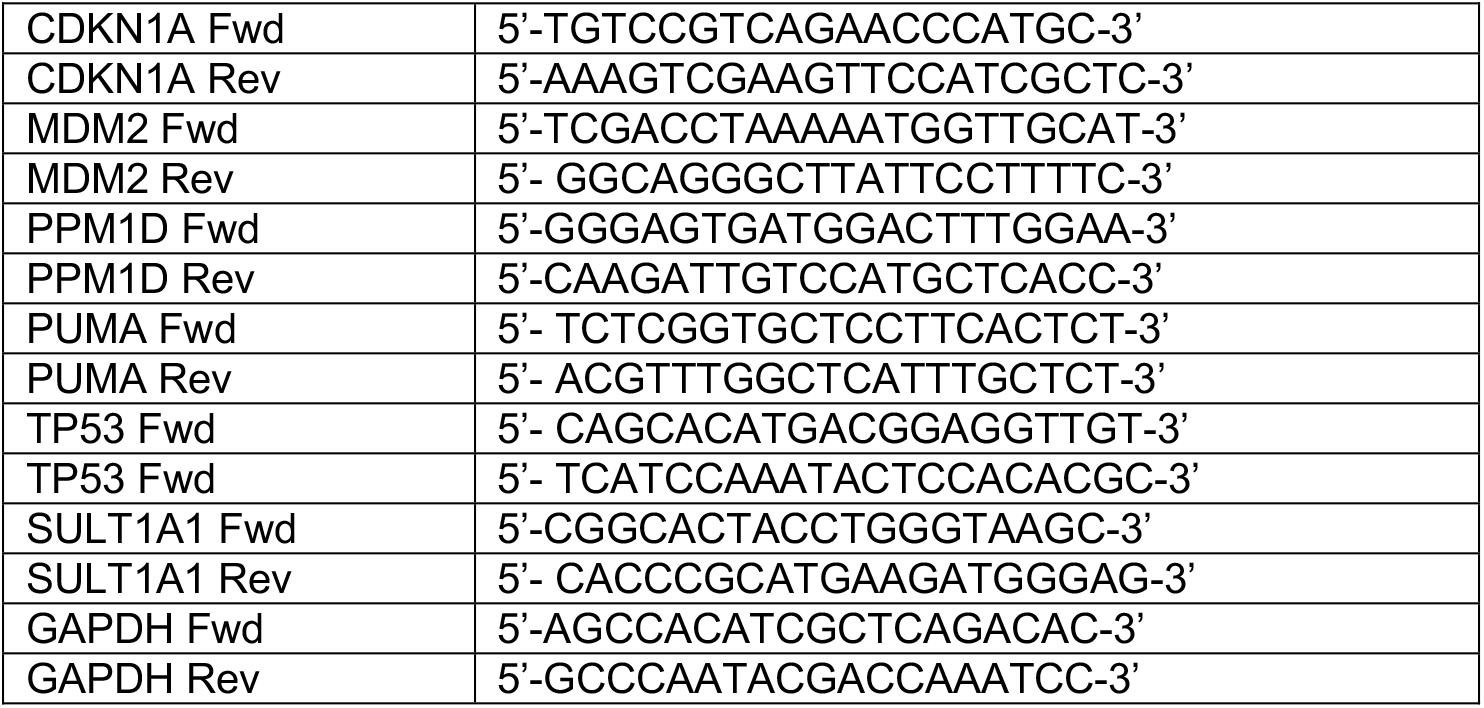

### Histone Extraction

This protocol was adapted with modifications from Dr. Junjie Chen’s laboratory (46) by Dr. Raul Mostoslavsky’s group. In brief, cells were lysed in buffer containing 10 mM HEPES pH 7.4, 10 mM KCl, and 0.05% NP-40 supplemented with protease and phosphatase inhibitors for 20 minutes on ice. The lysate was centrifuged for 10 minutes at 17,000 *g* at 4°C. The supernatant, which contains the cytoplasmic proteins, was removed and stored separately. The pellet, which contains the nuclear fraction, was washed once with lysis buffer and centrifuged for 10 minutes at 17,000 *g* at 4°C. The supernatant was discarded and the pellet was resuspended in 0.2 N HCl and incubated for 20 minutes on ice. The solution was centrifuged for 10 minutes at 17,000 *g* at 4°C. The supernatant, which contains histone proteins, was transferred to a fresh tube and protein concentration was measured using the Bradford assay.

### Western Blotting

For preparation of whole cell lysate, cells were lysed in lysis buffer (50 mM Tris-HCl pH 7.5, 150 mM NaCl, 1% Triton X-100, supplemented with protease and phosphatase inhibitors) for 30 minutes on ice. The lysate was centrifuged for 10 minutes at 17,000 *g* at 4°C to pellet cellular debris. The supernatant was transferred to a fresh tube and protein concentration was measured using the Bradford assay. 25 μg of protein was mixed with the NuPAGE LDS loading buffer (Life Technologies) containing the reducing agent dithiothreitol (DTT) and denatured by boiling at 95°C for 10 minutes. SDS-PAGE and membrane transfer was performed according to the manufacturer’s instructions (Life Technologies). The membrane was blocked in blocking buffer (PBS + 0.1% Tween-20 + 5% skim milk) for 1 hour at room temperature before incubating with the primary antibody diluted in the blocking buffer at 4°C overnight. After washing three times with PBS-T (PBS + 0.1% Tween-20), the membrane was then incubated with a horseradish peroxidase (HRP)-conjugated secondary antibody diluted in blocking buffer for 1 hour at room temperature. After washing three times with PBS-T, the proteins on the membrane were visualised with the ECL chemiluminescence substrate (Thermo Scientific) per the manufacturer’s instructions.

Primary antibodies used for Western blotting are listed here: H3K36me3 (Abcam), histone H3 (Abcam), RRM2 (Santa Cruz Biotechnology), caspase-3 (Abcam), caspase-3 (Abcam), p53 (Santa Cruz Biotechnology), phospho-p53 Ser15 (Cell Signaling Technology), phospho-CHK1 (Ser317) (Cell Signaling Technology), CHK1 (Cell Signaling Technology), phospho-CHK2 (Thr68) (Cell Signaling Technology), CHK2 (Cell Signaling Technology), p21 (Cell Signaling Technology), SULT1A1 (R&D Systems), and GAPDH (Novus).

### Cell Cycle Analysis by BrdU Incorporation

10^5^ cells/well in 2 mL of complete DMEM were seeded in 6-well plates overnight at 37°C. Immediately prior to harvesting, cells were incubated in media containing 20 μM bromodeoxyuridine (BrdU) for 30 minutes at 37°C while protected from light. Cells were collected by trypsinisation and fixed in ice-cold 70% ethanol for at least 30 minutes. Cells were incubated for 20 minutes at room temperature in 2 M hydrochloric acid (which denatures double-stranded DNA). Cells were washed once in PBS, followed by once in PBS + 0.5% Tween-20 + 0.5% FBS. Cells were then incubated in blocking buffer (PBS with 2% FBS) containing an anti-BrdU monoclonal antibody (BD Biosciences) (1:100 dilution) for 90 minutes at room temperature or overnight at 4°C. Cells were washed twice with PBS and then incubated in blocking buffer containing the Alexa Fluor 488 secondary antibody (Life Technologies) (1:200 dilution) for 60 minutes at room temperature in the dark. Cells were then washed in PBS, resuspended in PBS + 0.1 mg/ml propidium iodide, and analysed using the FACSCalibur flow cytometer (Becton Dickinson).

### Immunofluorescence Staining of Foci

10^5^ cells/well in 2 mL of complete DMEM were seeded onto glass coverslips placed in 6-well plates overnight. At the desired time point after treatment, pre-extraction buffer (10 mM PIPES pH 6.8, 300 mM sucrose, 100 mM NaCl, 1.5 mM MgCl_2_, and 0.5% Triton X-100) was added for 2 minutes on ice. The cells were then fixed in 4% paraformaldehyde in PBS for 10 minutes at room temperature. Coverslips were washed 3 times in PBS and incubated in blocking buffer (PBS + 0.1% Triton X-100 + 1% BSA + 1% FBS) for 1 hour. Coverslips were then incubated with primary antibody in PBS + 1% BSA + 1% FBS overnight at 4°C. Unbound primary antibody was removed by washing 3 times for 5 minutes in PBS at room temperature, followed by incubation with the indicated secondary antibody for 60 minutes at room temperature in the dark. Slides were then washed twice for 5 minutes in PBS and once in PBS with Hoechst 33342 (1:1000, Molecular Probes) before mounting with VECTASHIELD® Antifade Mounting Media (Vector Laboratories). Slides were imaged using the Zeiss 710 confocal microscope with a 63X oil objective.

The primary antibodies used for immunofluorescence are: phospho-RPA (Ser33) (Bethyl Laboratories), 53BP1 (GeneTex), and γH2AX (Novus).

### DNA Fibre Assay

Cells were seeded overnight such that they reached approximately 20% confluency the following day. Cells were labelled with 30 mM CldU (Sigma-Aldrich) for 30 min, washed twice with PBS, and labelled with 250 mM IdU in the presence of 150 nM RITA or DMSO for another 30 min. Following pulse labelling, cells were quickly trypsinized and resuspended in PBS at 2.5 × 10^5^ cells/ml. Labelled cells were diluted 1:8 with unlabelled cells, and 2.5 μl of cells were mixed with 7.5 μl of lysis buffer (200 mM Tris-HCl, pH 7.5, 50 mM EDTA, and 0.5% (w/v) SDS) on a glass slide. After 9 min, the slides were tilted at 15°–45°, and the resulting DNA spreads were air-dried and fixed in 3:1 methanol/acetic acid overnight at 4°C. The DNA fibers were denatured with 2.5 M HCl for 1 hr, washed with PBS, and blocked with 2% (w/v) BSA in PBS + 0.2% Tween- 20 for 40 min. The CldU and IdU tracts were labelled (for 2.5 hr in the dark, at RT) with anti-BrdU antibodies recognizing CldU (rat; Abcam) and IdU (mouse; BD), respectively. After washing for 5 × 3 min in 0.2% (v/v) PBS-T, the following secondary antibodies were used (1 hr incubation, in the dark, at RT): anti–mouse Alexa Fluor 488 (Molecular Probes) and anti–rat Cy3 (Jackson ImmunoResearch Laboratories, Inc.). After washing for 5 × 3 min in PBS-T (0.2% (v/v)), the slides were air dried completely, mounted with 20 ml/slide ProLong® Gold Antifade Mountant (Life Technologies), and sealed to a coverslip with transparent nail polish. Microscopy was performed with a fluorescence microscope (Nikon Ni-E; 100X oil objective) and the images were processed with the Fiji software. At least 200 fibres were measured per condition.

### Modified Single-Cell Gel Electrophoresis (Comet) Assay

This assay was adapted from the published protocol of Spanswick and colleagues (47). Briefly, cells were seeded at 100,000 cells/well in 6-well plates and incubated overnight. The following day, the cells were treated with DMSO, RITA, or mitomycin C for 1 hour at 37°C. All samples were then harvested by trypsinisation and aliquoted into separate Eppendorf tubes with 20,000-25,000 cells each, resuspended in freezing medium (complete DMEM + 10% DMSO), and frozen at −80°C for at least 24 hours prior to Comet analysis.

On the day of Comet analysis, the samples were thawed and resuspended in complete culture medium. With the exception of the unirradiated controls, the samples were irradiated on ice at 5 Gy (Xstrahl RS320 X-Ray Irradiator system). The cells were then pelleted by centrifugation and the remainder of the assay was performed under red light. Cells were placed on ice before being resuspended in 190 μL molten 0.6% LMP agarose cooled to 37°C. Then, 80 μL of the cell/LMP agarose mix was pipetted onto the centre of a dried 1% NMP agarose pre-coated slide, a coverslip placed on top and the slide then placed on ice to allow the gel to set. After the agarose had set, the coverslips were removed and the slides were placed in a tray on ice. The slides were covered with ice-cold lysis buffer containing 1% Triton X-100 and stored over night at 3°C. The slides were then removed from lysis buffer and washed with ice-cold double-distilled water, followed by incubation for 15 minutes in the dark. This step was repeated a further two times before the slides were carefully removed and transferred to an electrophoresis tank. The slides were covered with ice-cold alkali buffer and incubated for 20 minutes. Electrophoresis was performed for 20 minutes at 30 V (0.6 V/cm), 300 mA. The slides were then carefully removed from the buffer and placed on a horizontal slide rack. Each slide was flooded twice with 1 ml neutralisation buffer and incubated for 10 minutes, followed by rinsing twice with 1 mL PBS for 10 minutes. Excess liquid was removed, the slides were dried overnight in an incubator at 37°C.

To stain the slides for comet visualisation, the slides were re-hydrated in double-distilled water for 30 minutes. Each slide was then flooded twice with 1 ml of 2.5 μg/ml propidium iodide solution and incubated for at least 30 minutes at room temperature in the dark. The slides were rinsed twice with double-distilled water twice for 10 minutes 37°C in the dark. Once dry, a few drops of distilled water were added to each slide and covered with a coverslip. Comets were visualised at X200 magnification using an Olympus BH-2-RFL-T2 fluorescent microscope fitted with an excitation filter of 515-535 nm and a 590 nm barrier filter, and images were captured via a high performance CCDC camera (COHU MOD 4912-5000/0000). % tail DNA was calculated using Komet software (Andor Technology), with 100 comets scored per slide.

## Acknowledgments

We would like to thank Valentine Macaulay, Benoit Van den Eynde, and Ross Chapman for generously providing cancer cell lines, as well as Kilian Huber and Peter McHugh for useful discussions. This research was supported by the Medical Research Council, Cancer Research UK, and the Oxford-based charity UCARE (Urology Cancer Research and Education).

