## Supplementary Figures 1-7 for "Therapeutic targeting of SETD2-deficient cancer cells with the small-molecule compound RITA"

**This PDF file includes:**

Supplementary Table 1
Supplementary Figures 1 to 7

**Supplementary Data**

**Table S1.** List of compounds in the Oncology Drug Library (ODL) and Selleck Library that were included in the high-throughput screen.

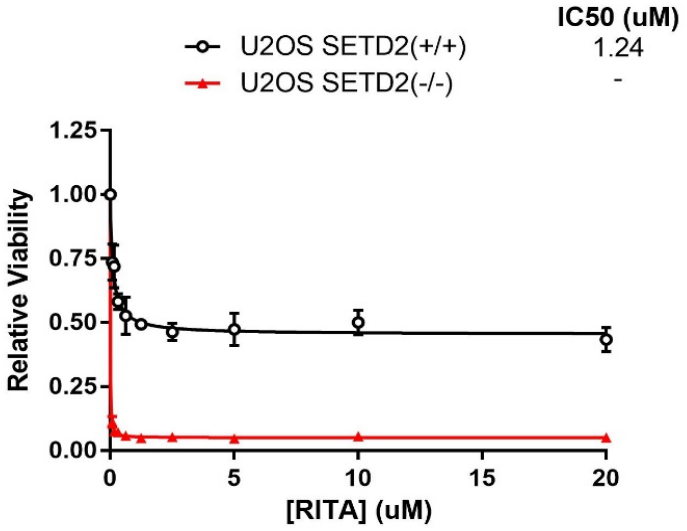


**Fig. S1.** Dose-response viability curve for RITA in parental and SETD2-CRISPR U2OS cells. Data are shown as mean ± SD (n ≥ 4). IC50 values were calculated via nonlinear regression (4-parameter curve fitting) when possible.

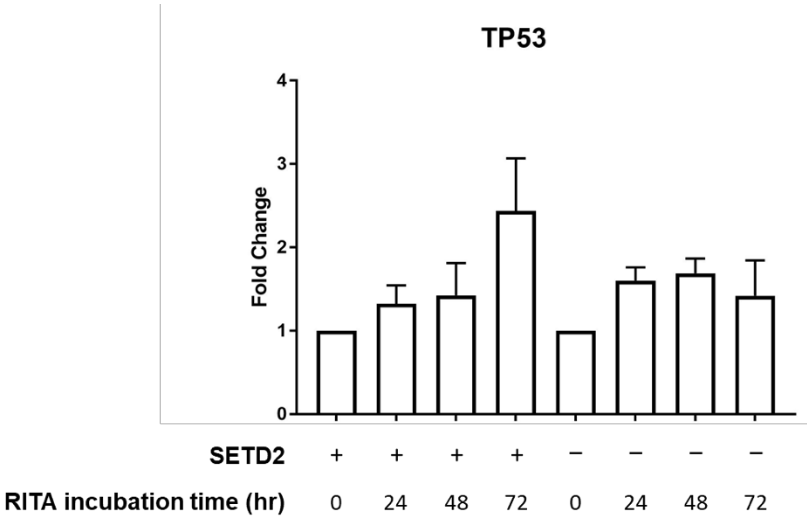


**Fig. S2.** Quantitative RT-PCR analysis of TP53 gene expression after RITA treatment. Samples were normalised to the housekeeping gene GAPDH. Fold change was calculated relative to untreated controls. Data are shown as mean ± SD.

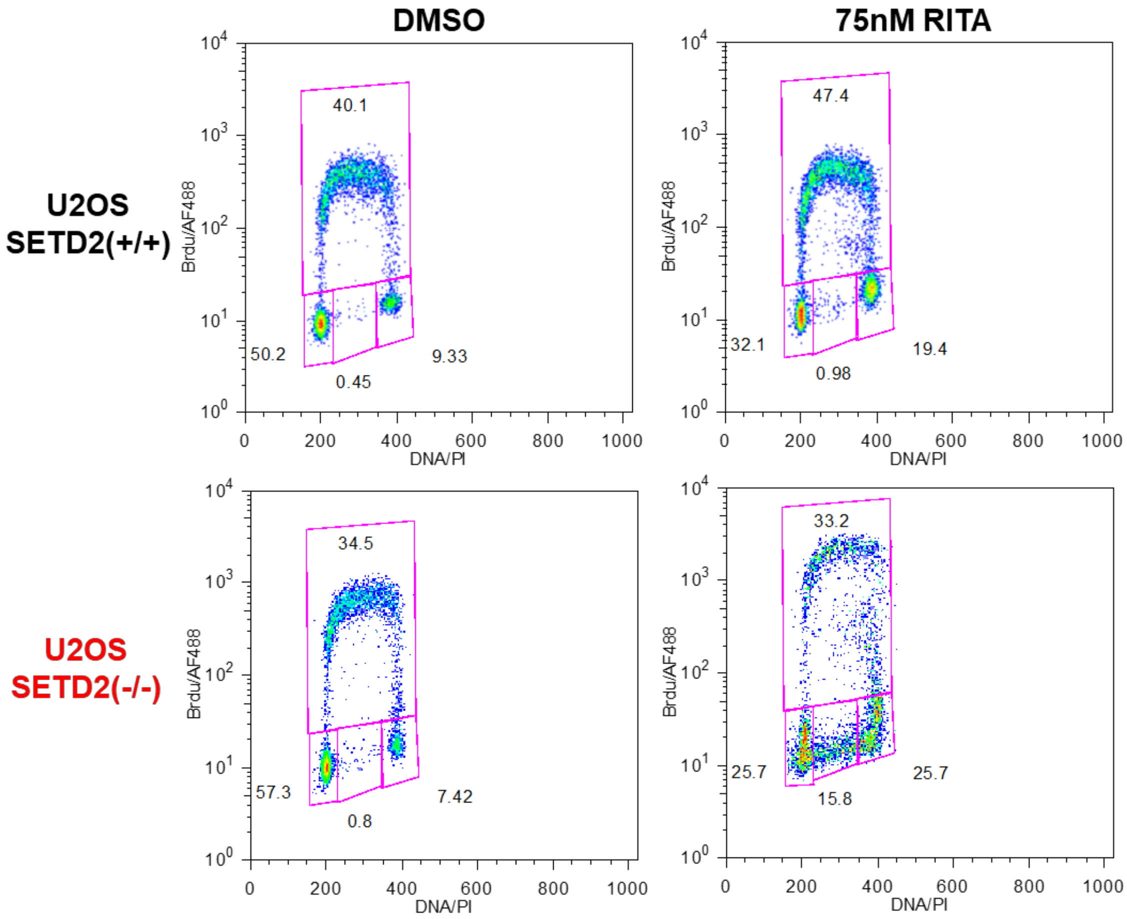


**Fig. S3.** BrdU/PI cell cycle profiles of U2OS cells after DMSO or RITA treatment. Images are representative of 3 independent experiments.

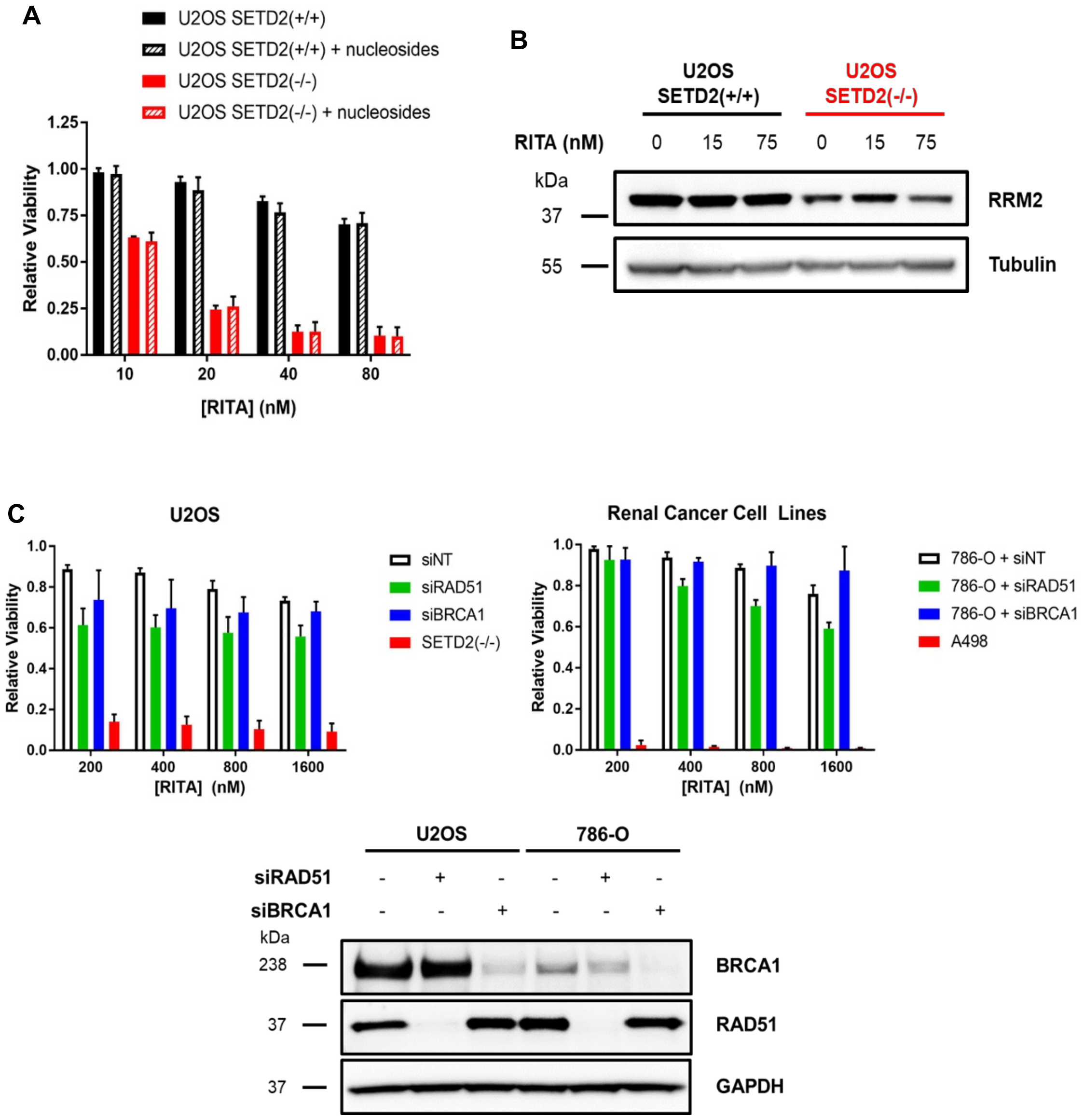


**Fig. S4.** RITA sensitivity of SETD2-deficient cells is not associated with nucleotide pools or homologous recombination. **(A)** Cell viability assay after RITA treatment of U2OS cells in the presence or absence of exogenous nucleosides. Data are shown as mean ± SD (n = 3). **(B)** Western blot of RRM2 protein in parental and SETD2-CRISPR U2OS cells treated with the indicated doses of RITA for 24 hours. **(C)** Cell viability assay after RITA treatment of U2OS cells or 786-O cells transfected with non-targeting (NT), RAD51, or BRCA1 siRNA. Data are shown as mean ± SD (n = 3). siRNA knockdown efficiency was confirmed by Western blot.

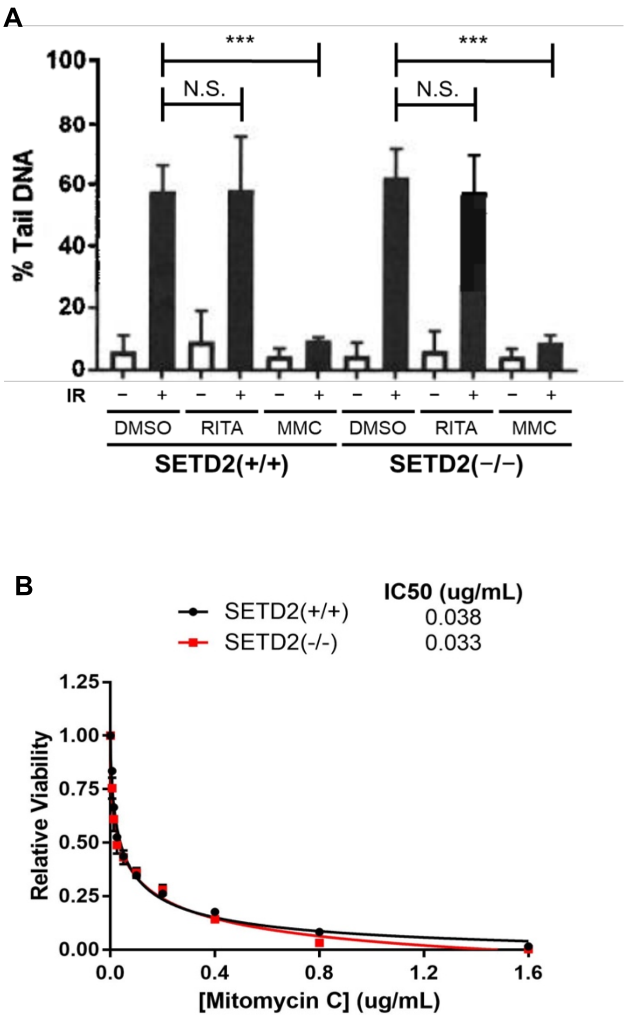


**Fig. S5.** RITA sensitivity in the absence of SETD2 is not associated with DNA crosslink formation. **(A)** Amount of fragmented (tail) DNA after ionising radiation (IR) as measured by comet assay in U2OS cells treated with RITA or mitomycin C (MMC). Data are shown as mean ± SEM (n = 3), each with 3 slides where 100 comets were counted per slide. P-values were calculated using the Student's two-tailed t-test. *** $p < 0.001$, N.S. = non-significant. **(B)** Dose-response viability curves for mitomycin C in U2OS cells. Data are shown as mean ± SD (n = 3). IC50 values were calculated via nonlinear regression (4-parameter curve fitting).

A

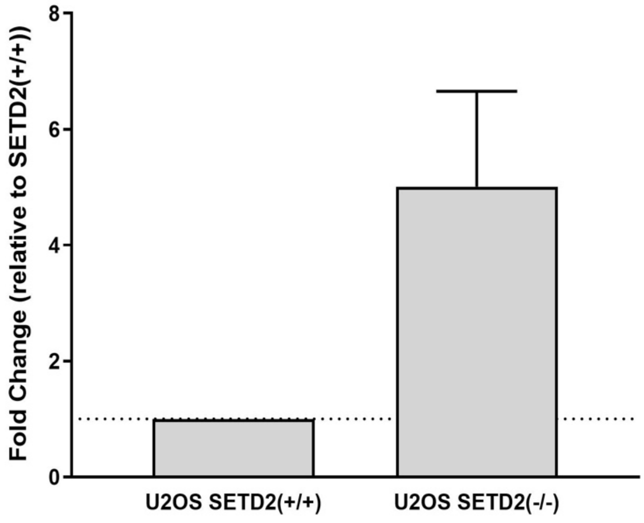


B

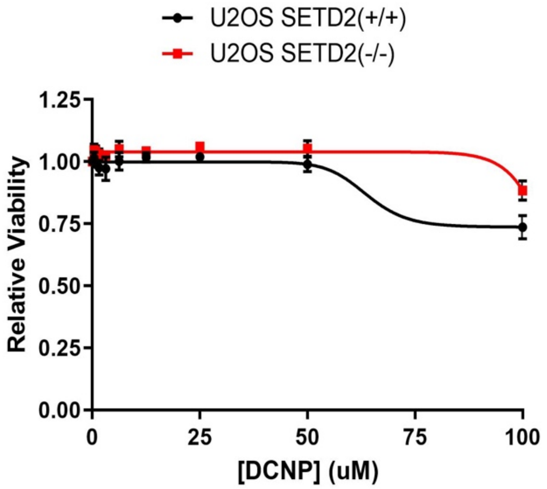


**Fig. S6.** RITA sensitivity in the context of SETD2 loss is correlated with expression levels of the phenol sulphotransferase SULT1A1. **(A)** Quantitative RT-PCR analysis of *SULT1A1* gene expression in U2OS cells. Samples were normalised to the housekeeping gene *GAPDH*. Fold change was calculated relative to wild-type SETD2(+/+) U2OS. Data are shown as mean ± SD. **(B)** Dose response viability curves for U2OS cells treated with the phenol sulphotransferase inhibitor DCNP. Data are shown as mean ± SD.

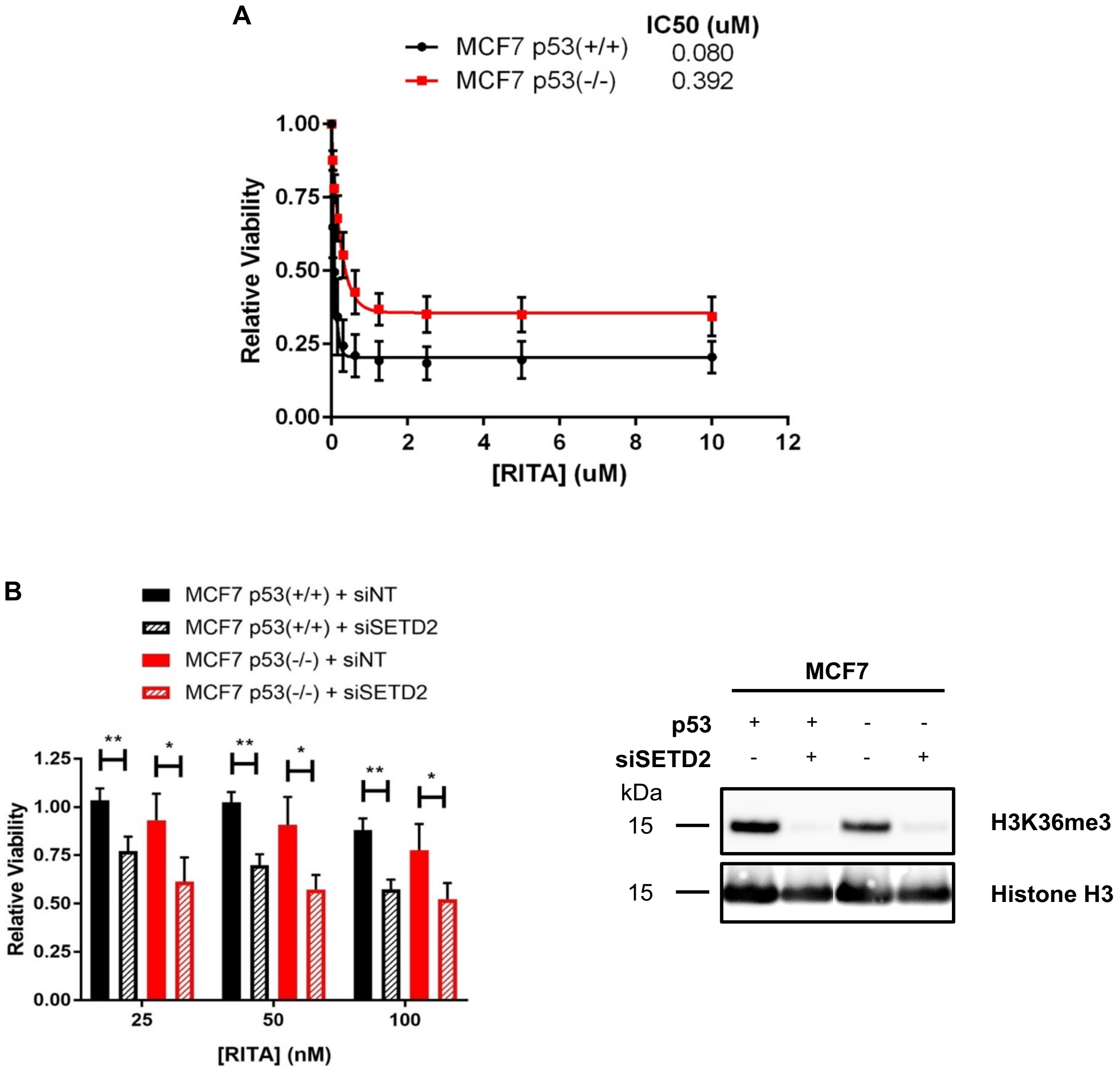


**Fig. S7.** RITA is more potent in the presence of p53 but SETD2 loss sensitises cells to RITA regardless of p53 status. **(A)** Dose-response viability curves for RITA in MCF7 cells. Data are shown as mean ± SD (n = 3). IC50 values were calculated via nonlinear regression (4-parameter curve fitting). (**B)** Cell viability assay after RITA treatment combined with siRNA depletion of SETD2 in MCF7 cells. Viability was calculated relative to cells incubated with non-targeting siRNA. Data are shown as mean ± SD (n = 3). P-values were calculated using the Student's two-tailed t-test (* $p < 0.05$, ** $p < 0.01$). SETD2 knockdown efficiency was confirmed by Western blotting for H3K36me3.
